# Sleep differentially affects early and late neuronal responses to sounds in auditory and perirhinal cortices

**DOI:** 10.1101/743666

**Authors:** Y. Sela, AJ. Krom, L. Bergman, N. Regev, Y. Nir

## Abstract

A fundamental feature of sleep is reduced behavioral responsiveness to external events, but the extent of processing along sensory pathways remains poorly understood. While responses are comparable across wakefulness and sleep in auditory cortex (AC), neuronal activity in downstream regions remains unknown. Here we recorded spiking activity in 435 neuronal clusters evoked by acoustic stimuli in the perirhinal cortex (PRC) and in AC of freely behaving male rats across wakefulness and sleep. Neuronal responses in AC showed modest (around 10%) differences in response gain across vigilance states, replicating previous studies. By contrast, PRC neuronal responses were robustly attenuated by 47% and 36% during NREM sleep and REM sleep, respectively. Beyond the separation according to cortical region, response latency in each neuronal cluster was correlated with the degree of NREM sleep attenuation, such that late (>40ms) responses in all monitored regions diminished during NREM sleep. The robust attenuation of late responses prevalent in PRC represents a novel neural correlate of sensory disconnection during sleep, opening new avenues for investigating the mediating mechanisms.

**Significance Statement:** Reduced behavioral responsiveness to sensory stimulation is at the core of sleep’s definition, but it is still unclear how the sleeping brain responds differently to sensory stimuli. In the current study we recorded neuronal spiking responses to sounds along the cortical processing hierarchy of rats during wakefulness and natural sleep. Responses in auditory cortex only showed modest changes during sleep, whereas sleep robustly attenuated the responses of neurons in high-level perirhinal cortex. We also found that during NREM sleep, the response latency predicts the degree of sleep attenuation in individual neurons above and beyond their anatomical location. These results provide anatomical and temporal signatures of sensory disconnection during sleep and pave the way to understanding the underlying mechanisms.

## Introduction

Reduced behavioral responses to external sensory stimuli is an essential definition of sleep (Carskadon and Dement, 2011), and is ubiquitous across the animal kingdom (Cirelli and Tononi, 2008). Yet, how sleep affects the responses along sensory pathways remains poorly understood. During sleep, external stimuli rarely affect perception (Nir and Tononi, 2010), but it is also clear that some discriminative sensory processing persists (Andrillon et al., 2016). Surprisingly, only few studies investigated sensory responses during natural sleep at the neuronal level (Hennevin et al., 2007). Early studies (Gücer, 1979; Livingstone and Hubel, 1981) contributed to the “thalamic gating” model (Steriade and McCarley, 1990), proposing that thalamic burst-mode activity and sleep spindle oscillations disrupt signal propagation to the cerebral cortex during sleep (McCormick and Bal, 1994; Steriade, 2003). However, this view was challenged by sleep attenuation of responses in the olfactory cortex that are not relayed through thalamus (Murakami et al., 2005). In addition, several recent studies in the auditory system showed that responses in the primary and secondary auditory cortex are comparable across wakefulness and natural sleep (Pena et al., 1999; Edeline et al., 2001; Issa and Wang, 2008; Nir et al., 2013), even at times when sleep spindles occur (Sela et al., 2016).

Do sounds effectively modulate responses in downstream association cortex during sleep? On one hand, hippocampal-dependent memory consolidation can be promoted via Targeted Memory Reactivation (TMR) that re-presents sounds that were used as training cues or context during sleep (Oudiette and Paller, 2013). TMR has been shown to influence hippocampal activity during sleep (Rasch et al., 2007; Rothschild et al., 2017), suggesting that sounds can modulate activity in association cortices. On the other hand, non-invasive imaging in humans suggests that during sleep, auditory responses downstream from auditory cortex (AC) are attenuated: studies using magnetoencephalography (MEG) (Kakigi et al., 2003; Strauss et al., 2015), electroencephalography (EEG) (Makov et al., 2017), and functional magnetic resonance imaging (fMRI) (Portas et al., 2000) suggest comparable responses across vigilance states in early sensory regions and robust attenuation in association cortex. However, investigation of sensory responses downstream from sensory cortical regions at the level of neuronal spiking activity is still missing.

We hypothesized that the perirhinal cortex (PRC) constitutes a high-order cortical region that may show robust attenuation of sensory responses during sleep. The perirhinal cortex is a multi-modal region downstream to several primary sensory cortices, where information converges from different sensory modalities to support higher brain functions including perception and recognition memory (Kealy and Commins, 2011). Moreover, it has been shown to contain intrinsic inhibitory mechanisms that affect information flow to the hippocampus (Decurtis and Pare, 2004; Pelletier, 2004). Perirhinal cortex is located further downstream from the suprarhinal auditory field (SRAF) (Profant et al., 2013; Lee et al., 2016). While perirhinal neurons reliably respond to sounds during wakefulness (Allen et al., 2007; Furtak et al., 2007), separate studies during anesthesia did not reveal significant auditory responses as one moves ventrally towards rhinal fissure (Doron et al., 2002). For the above reasons, we set out to directly compare auditory responses in perirhinal cortex neurons across wakefulness and natural sleep. We also compared perirhinal response profiles to simultaneously recorded neurons in auditory cortex. We find that perirhinal cortex neurons robustly attenuate their auditory responses during sleep, in contrast to auditory cortex neurons that are minimally affected by vigilance state. In addition, we find that the response latency of each neuronal cluster (typically <20ms in AC and >40ms in PRC) is correlated with the degree of sleep attenuation, and that in NREM sleep latency better predicts sleep attenuation than does cortical region.

## Methods

### Animals

Experiments were performed in 8 Long-Evans and 2 Wistar adult male rats (250-350 g) individually housed in transparent Perspex cages with food and water available ad libitum. Ambient temperature was kept at 20-24° Celsius, and light-dark cycles of 12/12h at 10AM/PM. All experimental procedures including animal handling, surgery, and experiments followed the NIH Guide for the Care and Use of Laboratory Animals and were approved by the Institutional Animal Care and Use Committee (IACUC) of Tel Aviv University.

### Surgery and implantation

Animals were anesthetized using Isoflurane (4-5% induction, 1-2% maintenance) and placed in a stereotactic frame (David Kopf Instruments; CA, USA), while maintaining body temperature around 37<u>°</u> Celsius using a closed-loop heating pad system (Harvard Apparatus, MA, USA). The head was shaved and Viscotear gel was applied to protect the eyes. After exposing and cleaning the skull, frontal and parietal screws (1mm in diameter) were placed over the left hemisphere for EEG recording. Two screws were placed above the cerebellum as reference and ground, and an additional anchor screw above the right frontal lobe for stabilization. Two single stranded stainless steel wires were inserted to either side of neck muscles to measure EMG (Figure 1A). EEG and EMG wires were soldered onto a headstage connector (Omnetics Connector Corporation; MN, USA). Dental cement was used to cover all screws and EEG/EMG wires.

**Figure 1.**
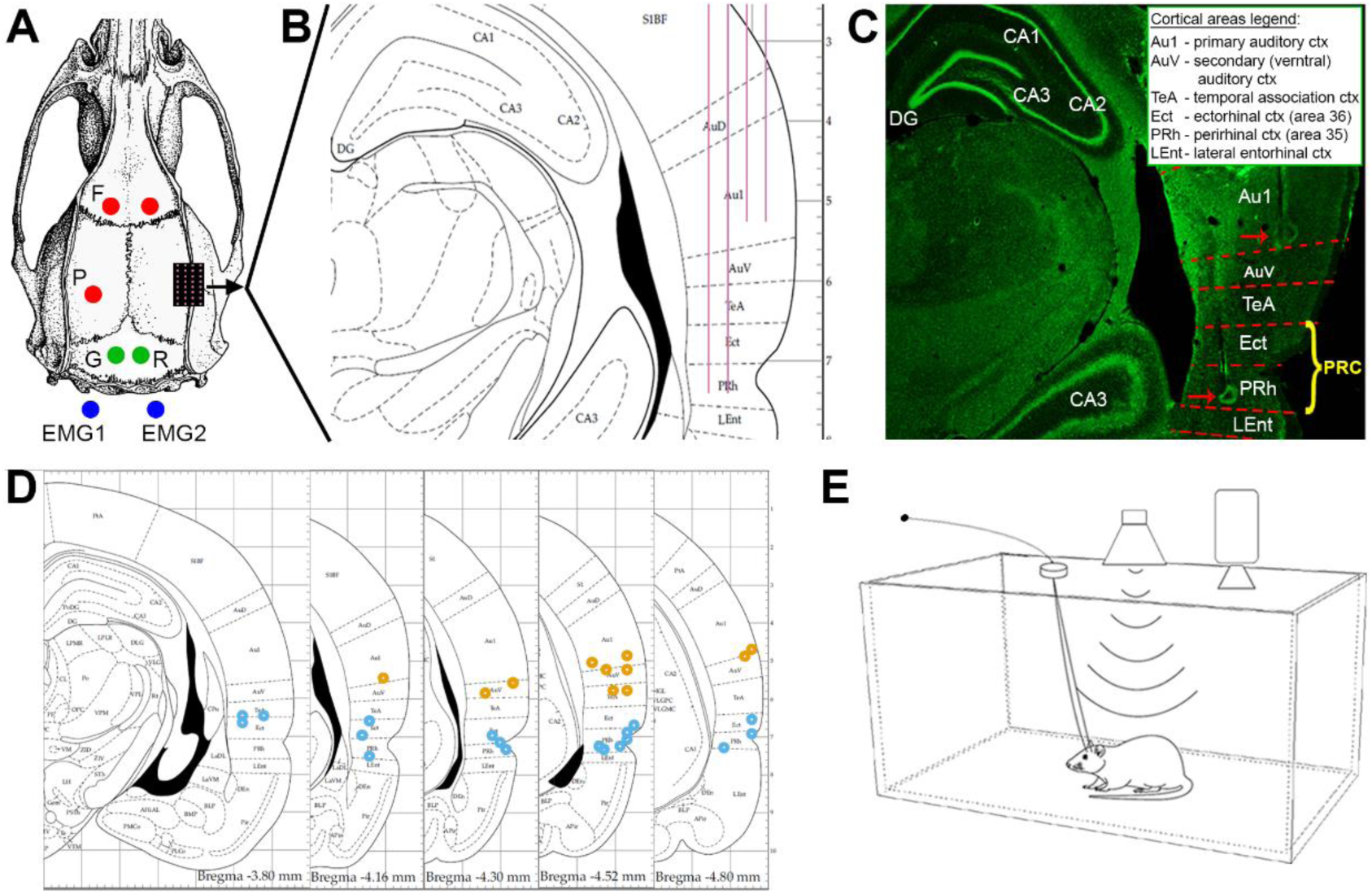
Experimental Design. (A) Surgical plan, including frontal (“F”) and parietal (“P”) EEG screws (red circles), ground (“G”) and reference (“R”) screws (green circles) above cerebellum, and two wires (blue circles) recording EMG from neck muscles. Microwire arrays were implanted in rat temporal lobe through a craniotomy (black rectangle with purple dots marking each wire). (B) Coronal section (−4.16mm from Bregma) of the implantation area, showing the implantation plan of 32-channel microwire arrays (4 rows × 8 wires each) targeting AC (2 shorter rows) and PRC (2 longer rows). (C) Representative histological verification of electrode positions. Red dashed lines denote estimated area borders. Red arrows point to detectable electrode tips in AC and PRC. Abbreviations of brain regions in inset. PRC (yellow font) includes both area 36 (Ectorhinal cortex (Kealy and Commins, 2011)) and area 35 (PRh) (D) Five coronal sections (spanning -3.80 to -4.80 from Bregma) showing the locations of electrodes identified in histology (orange/blue circles for AC/PRC microwires). (E) Experimental setup diagram: rats were placed individually in a sound-attenuating chamber, while auditory stimuli were delivered free-field from a speaker mounted 55cm above the cage center. Intracranial electrophysiology and sleep polysomnography (EEG, EMG, video) were continuously collected while rats were behaving and sleeping freely.

**Figure 2.**
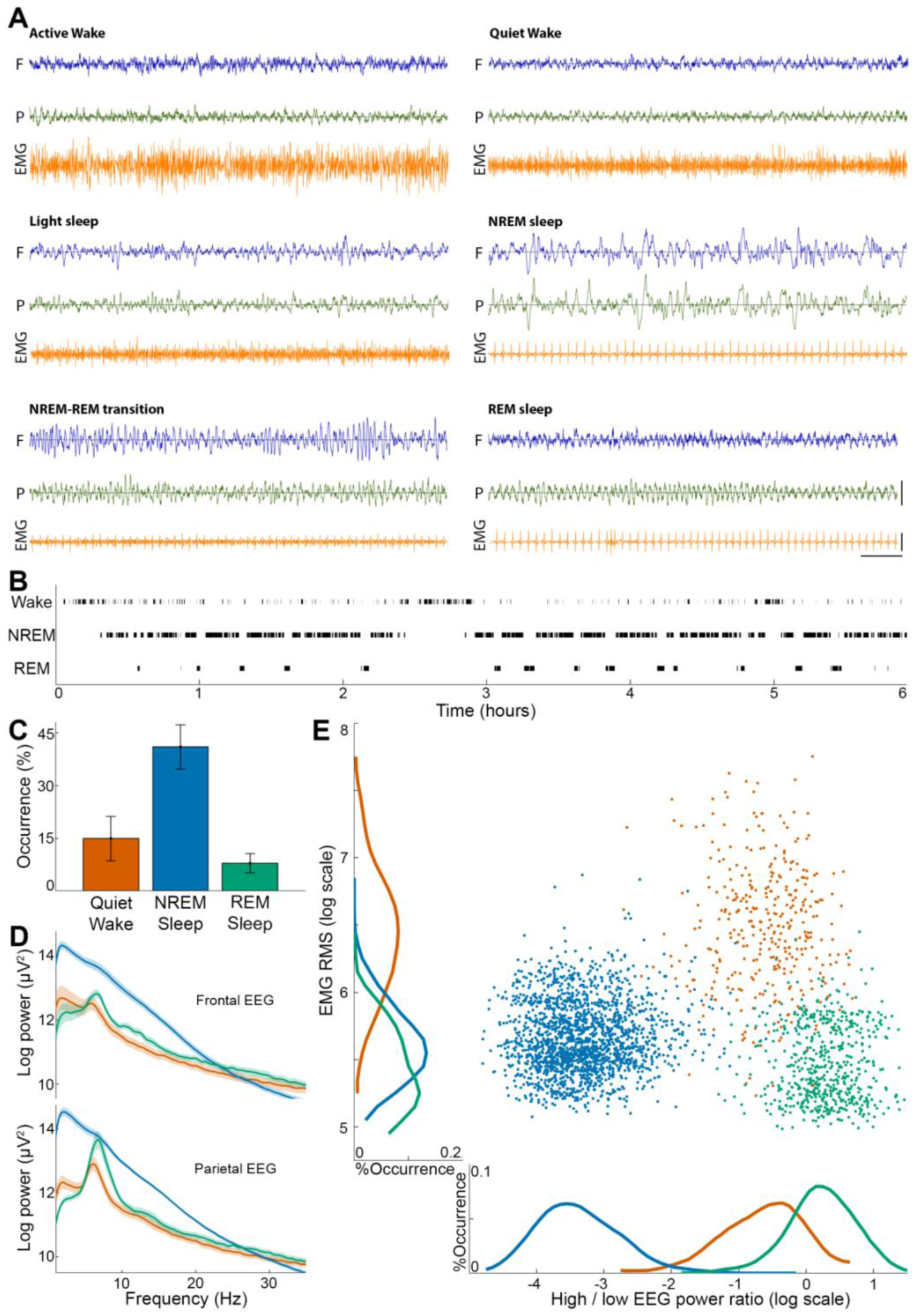
Sleep scoring. (A) Representative polygraphic traces of the different vigilance states. In each panel, electroencephalogram (EEG) above the frontal (blue) and parietal (green) cortices are shown above the electromyogram (EMG, orange). Horizontal line at bottom right corner indicates 1 second time scale, and vertical lines represent 1mV and 0.25mV for EEGs and EMG respectively. (B) Hypnogram showing the time-course of vigilance state transitions in a representative experiment. Black ticks denote the states used for full analysis (top row, quiet wake; middle row, NREM sleep; bottom row, REM sleep). (C) Occurrence (percent of time, y-axis) of vigilance states (x-axis) during experiments. Error-bars mark SD across sessions. Colors in all panels represent vigilance states (red, quiet wake; dark blue, NREM sleep; green, REM sleep). (D) EEG spectral power across vigilance states in frontal EEG (top) and parietal EEG (bottom). Thick lines and shading represent mean ± SEM across experimental sessions. (E) A representative scatter plot distribution of EMG RMS (y-axis) vs. frontal EEG power distribution (ratio between power in high (>25 Hz) vs. low (<5 Hz) frequencies, x-axis) in each 4s epoch. Note that both sleep states have low EMG, while EEG is dominated by high-frequency activity in wake and REM sleep.

A small craniotomy was performed over the right hemisphere and the dura was reflected under microscopic control. Multi-channel microwire arrays (each wire: 50 µm diameter, 45° tip angle) were implanted (Tucker-Davis Technologies Inc (TDT), Alachua, FL, USA). The arrays consisted of either 16 wires (2 rows * 8 channels, 9.5mm long) or 32 wires (4 rows * 8 channels, two short 7.5mm rows and two long 9.5mm rows), with 375µm medial-lateral separation between rows and 250µm anterior-posterior separation within each row. Implantation was made either diagonally (insertion point P: 3.6 mm, L: 4.2 mm relative to Bregma and inserted to a depth of 8mm) in angle of 22° (n=3) or vertically (P: 3.6 mm, L: 6.65 mm, D: 7.4 mm relative to Bregma) after the temporalis muscle was gently separated and retracted from the bone (n=7, Figure 1B). After implantation, the craniotomy was covered with silicon gel (Kwik-Sil; World Precision Instruments, FL, USA) and the electrode array was fixed in place with Fusio (Pentron, Czech Republic). Dexamethasone (1.3 mg/kg) was given during recovery with food to reduce pain and edema around implanted electrodes.

### Histology

At the end of each experiment, rats were deeply anesthetized (5% isoflurane) and transcardially perfused with saline followed by 4% paraformaldehyde. Brains were refrigerated in paraformaldehyde for at least a week, and sectioned in 50-60 µm serial coronal sections using a Vibrating Microtome (Leica Biosystems, Israel). Slices were stained with fluorescent cresyl-violet (Rhenium, Modi’in, Israel)

### Electrophysiology and spike sorting

All electrophysiological data were acquired using an RZ2 high performance processor (TDT). We continuously recorded extracellular spiking activity (24.4 kHz, filtered 300-5000 Hz) and local field potential (filtered 0.5-300 Hz) from the same microwire electrodes. Epidural EEG (filtered 0.5-300 Hz) and EMG (filtered 10-100 Hz) were first pre-amplified (RA16LI, TDT). All data were digitally sampled by a PZ2 amplifier (TDT) and synchronized with a video system RV2 (TDT). LFP, EEG and EMG data were resampled to 1000Hz using custom Matlab scripts (The MathWorks, Natick, MA, USA). Spike sorting was performed offline with ‘wave_clus’ (Quiroga et al., 2004) using a detection threshold of 5 SD above the median (median(|data|/0.6745, Figure 3A), and applying automatic superparamagnetic clustering of wavelet coefficients followed by manual refinement based on consistency of spike waveforms and inter-spike-interval distribution (Figure 3B&C).

**Figure 3.**
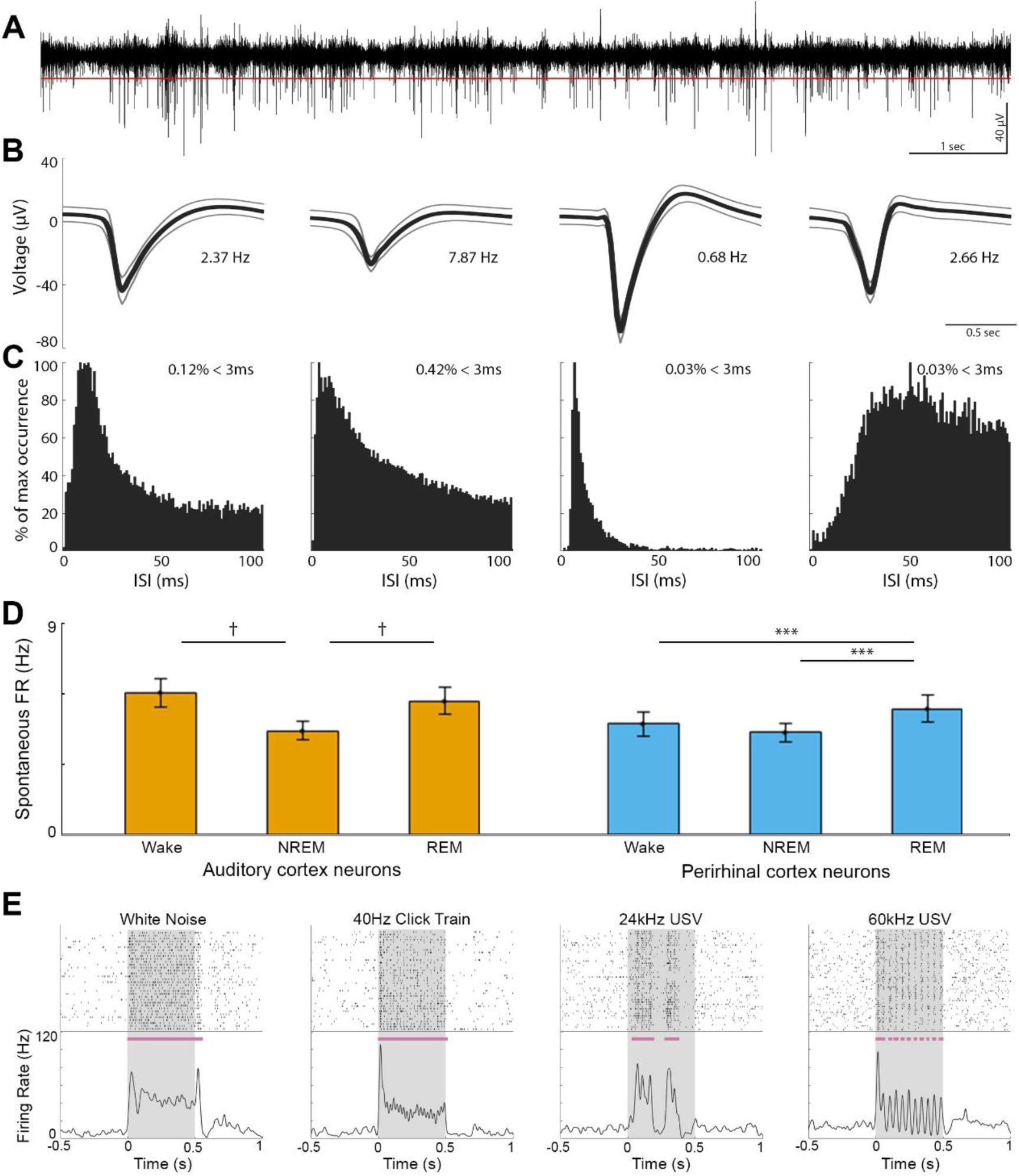
Neuronal spiking activity in AC and PRC. (A) Representative ten second trace of high-pass filtered (300-5000 Hz) local field potential used for identification of spiking activity. Red line denotes the automatic spike detection threshold. (B) Action potential waveforms of three detected single units and one multiunit cluster (the second from left) in the above channel. Thick lines and thinner frames mark mean ± SD. Numbers represent the mean firing rate of each unit. (C) Corresponding inter-spike-interval histograms of each unit. Percentage in upper right corner denotes the prevalence of action potentials occurring successively within 0 - 3ms. (D) Baseline firing rate across vigilance states in AC (left, orange, n=178) and PRC (right, blue, n=132) neurons. Error bars represent SEM across neurons. ***p = 0.001, †p = 10^−8^, via signed-rank test. (E) Representative raster plots (top) and peri-stimulus time histograms (PSTHs, bottom) of neuronal responses. Grey shading denotes stimulus presentation times. Horizontal purple bars denote automatically detected response intervals (Methods). USV, ultrasonic vocalization.

### Auditory stimulation

All experiments were conducted in a sound-attenuation (−55dB) chamber (H.N.A, Kiriat Malachi, Israel). Sounds were synthesized in Matlab, transduced to voltage signal (195kHz sampling rate, RZ6, TDT), passed through a zero-amplification speaker amplifier (SA1, TDT), and played free field via a magnetic ultrasonic speaker (MF1, TDT) mounted 55 cm above the center of the cage (Figure 1E). Given the free-field setup, stimulus intensity at the animal’s ears in each trial inevitably varies depending on precise position relative to speaker. We measured sound intensity (values below) with an ultrasonic calibration microphone (PCB Piezotronics model 378C01, Depew, NY, USA) placed directly below the speaker in the center of the cage. During experiments, a wide battery of 36 sounds (500ms each, except for a 1ms click stimulus) were delivered, including: pure tone pips (3kHz, 6kHz, 12kHz, 24kHz, 48kHz) at three different volumes (76, 82, 93 db SPL); single octave band-passed white noise (centered on the same frequencies as tone pips); 40Hz click trains (with 3 intensities as above), and single clicks. In addition, we presented behaviorally relevant sounds of pre-recorded (Avisoft Bioacoustics; www.avisoft.com) ultrasonic rodent vocalizations (USV, 24 kHz, 42 kHz, 66 kHz, and six types of 60kHz), were concatenated to produce 500ms-long stimuli. Finally, in an attempt to generate behaviorally-meaningless stimuli with near-identical physical features, three additional vocoded vocalizations were generated by taking the first three USVs above, identifying their principle frequency, and modulating a pure tone of this frequency using their original sound intensity envelopes.

### Experimental Design

Three days prior the surgery, rats were moved with their home cage inside an acoustic chamber for habituation to the experimental environment. After surgery, a recovery of one week was allowed undisturbed for renormalization of the sleep/wake. Next, 2-3 additional days were used for habituation to recording cable and to auditory stimulation. Experimental sessions lasted 5-9h and started between 10:00-12:00 to maximize REM sleep trials in the late ‘lights-on’ afternoon sleep hours. All 36 stimuli were presented in a pseudorandom order with an average inter-stimulus-interval of 2.5±0.4s (with jitter). There was a pause of 10 minutes after every cycle of 24 repetitions of all stimuli.

### Sleep Scoring

Sleep scoring was based on a combined manual examination of frontal and parietal EEG, EMG, and video data, using custom Matlab software, which enabled precise marking of state transitions. Data were divided to six vigilance states (Figure 2A) as in (Datta and Hobson, 2000): (1) Quiet wake: low voltage-high frequency EEG, and high tonic EMG. (2) Active wake: low voltage-high frequency EEG, accompanied by phasic EMG activity and behavioral activity (e.g. eating, grooming or exploring) confirmed with video. (3) NREM sleep: high-amplitude slow wave activity and low tonic EMG activity. (4) Light NREM sleep: transition state between wakefulness and NREM sleep, characterized by EEG slowing without obvious slow waves co-occurring with reduced EMG (5) REM sleep: high-frequency wake-like frontal EEG co-occurring with theta activity in parietal EEG and flat EMG. (6) NREM-REM transition: transition into REM sleep showing mixed EEG signatures of both NREM and REM sleep (e.g. weak theta in parietal EEG with spindle activity in frontal EEG). Subsequent analysis of auditory responses focused on quiet wakefulness, NREM sleep, and REM sleep and did not include periods of light NREM sleep (14±2.6%), NREM-REM transition periods (3±1.3% of data), active wake (13±4.8% of data), or intervals not matching these six states (5±2.8% of data). Joint-distributions of EMG levels and EEG high-/low-frequency power ratio (Figure 2E) were calculated based on 4s epochs. Power ratio was defined as power>25Hz divided by power<4Hz in the frontal EEG.

### Analysis of auditory responses

#### Detection of significant responses

We detected time intervals of significant increased neuronal activity in responses to auditory stimuli as follows: First, each trial was smoothed by convolution with a Gaussian kernel (σ=8ms). Then, a one tailed Mann-Whitney test compared, across trials, each millisecond within a 600ms interval (500ms of stimulus + 100ms following it), against all 500ms periods of baseline activity. We corrected for the 600 multiple comparisons using False-Discovery-Rate (Benjamini and Yekutieli, 2001) with base alpha of 0.001. Responses shorter than 12ms were excluded, and undetected intervals shorter than 4ms that preceded and followed responses were categorized as misses and bridged with adjacent intervals. In REM sleep, where less data were available, we restricted analysis to those stimuli that had at least 15 trials.

#### Latency of neuronal responses

To detect the peak latency of each neuronal cluster, we first quantified its responses to pure tones and click stimuli during wakefulness. For each stimulus, the PSTH was smoothed with a Gaussian kernel (σ=8ms). Peak latency was first defined for each stimulus separately as the peak of a significant response within the first 100ms. Then, peak latency was defined for each neuron as the average of its peak latencies across stimuli. Computing response latencies jointly across all states did not reveal significant changes to the main results.

#### Definition of early and late responding neurons

Based on the result in Figure 6B we identified two distinct populations of neurons that could be reliably separated according to their peak response latency, where 85% of neurons with responses <40ms were in AC, and 89% of neurons with responses >40ms were in PRC. We therefore defined either early- or late-responding neurons as those with peak latency of <20ms or >40ms respectively, leaving out neurons with peak latencies of 20-40ms from the analysis based on latency to avoid ambiguity. Precise cutoffs for early vs. late neurons (e.g. 15/20/25ms vs. 30/40/50ms) did not significantly affect the results.

In order to identify early- and late-responding neurons on the same microwires (Figure 6E&F) we focused on channels with at least two units exhibiting a difference >20ms in response latency, with the shorter latency < 20ms.

#### Comparison across vigilance states

For each unit, stimulus, and pair of states to be compared (e.g. wakefulness vs. NREM sleep) separately, we identified temporal intervals with significant responses in either state (as above). The response magnitude in both states was computed in these same intervals, after subtraction the pre-stimulus baseline activity in each state. Then, gain modulation factors between vigilance state pairs for each neuronal cluster were calculated as:

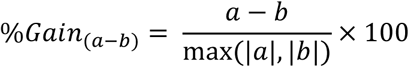

where a,b are the averaged response magnitudes in each state. Distributions of gain factors across neurons are summarized as median, and [lower and upper] 95% confidence intervals around the median (Campbell and Gardner, 1988).

#### Statistics

All data analysis was carried out using custom Matlab scripts. Unless stated otherwise, a two tail Mann-Whitney U test was used to assess statistical significance. Multiple-comparisons were corrected via False-Discovery-Rate (Benjamini and Yekutieli, 2001). Stimuli were presented in a pseudorandom order, and interleaved across vigilance states due to the natural fragmented sleep of rodents. Sleep scoring was blind to the timing of auditory trials.

## Data and code availability

The data that support the findings of this study and the computer code used to generate the results are available from the corresponding author upon reasonable request.

## Results

To study sensory processing along the auditory-mnemonic hierarchy across wakefulness and sleep, we implanted adult male rats (n=10) with microwires targeting the right AC (n=7) and/or PRC (n=7, Figure 1). After a week of recovery and habituation, we presented a wide battery of sounds (including tones, clicks and click trains, white noise, rat vocalizations, and vocoded vocalizations) in 12 experimental sessions lasting 5-8h each, as animals spontaneously switched between vigilance states in their home cage during the light phase. Sleep scoring was performed based on EEG, EMG and video (Figure 2A & Methods). Quiet wake, NREM sleep, and REM sleep composed 15±6.3%, 41±6.2%, and 8±2.4% of experiment time, respectively (mean±SD, Figure 2B&C) and exhibited characteristic changes in EEG power spectra including increased slow wave activity in NREM sleep, and higher theta/high-frequency activity in wakefulness and REM sleep (Figure 2D).

We isolated 435 neuronal clusters (130 single-units, 305 multi-unit clusters; Figure 3A,B&C) that significantly responded to auditory stimuli in at least one vigilance state (out of 485 neuronal clusters in total). To compare between activities in the two regions, neurons were separated into AC (n=178) and PRC (n=132) based on histology in all cases where that provided unequivocal information (n= 8 sessions in 6 rats) (Figure 1). Analysis of baseline firing rates revealed that AC neurons exhibited 21-26% higher firing rates during wakefulness (6.0±0.6Hz) or REM sleep (5.6±0.6Hz) compared to NREM sleep (4.4±0.4Hz, p values ≤ via signed-rank tests). In contrast, baseline firing rates of PRC neurons was higher during REM sleep (5.3±0.6Hz) in comparison to wakefulness (4.7±0.5Hz, 12% difference) and NREM sleep (4.3±0.4Hz, 23% difference; p-values≤0.001, via signed-rank tests, Figure 3D).

### Auditory responses in AC and PRC neurons across wakefulness and sleep

To examine auditory-evoked discharges, neuronal responses were aligned to sound onset, and an automatic algorithm detected significantly responsive intervals around stimulus presentation (Figure 3E & Methods). We compared auditory spiking responses across vigilance states in each cortical region. In line with previous studies (Issa and Wang, 2008; Nir et al., 2013), spiking responses in AC neurons exhibited similar profiles across vigilance states whereas PRC neurons exhibited weaker responses in both NREM and REM sleep (Figure 4).

**Figure 4.**
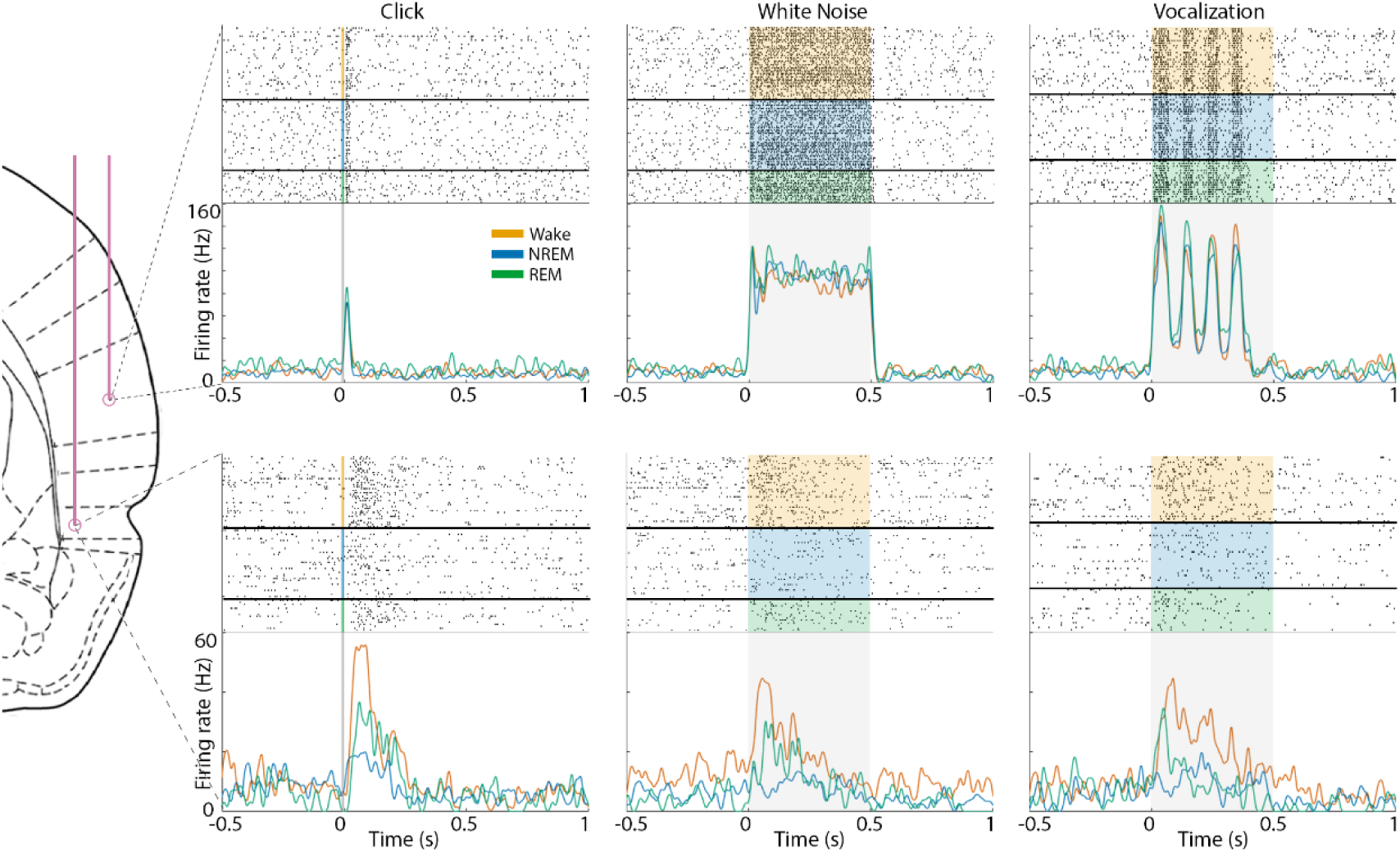
Example auditory responses in AC and PRC neurons across wakefulness and sleep. Raster plots (top panels) and PSTH (bottom panels) show representative auditory responses in an AC neuron (top row) and a PRC neuron (bottom row) recorded simultaneously. Columns show responses to different stimuli including a click (left), 12kHz white noise burst (middle), and 66kHz ultrasonic vocalization (right). In each panel shaded area represents stimulus duration, and raster plots are grouped by vigilance state (orange, wake; blue, NREM sleep; green, REM sleep). PSTH traces of different vigilance states are superimposed (colors as above).

We proceeded to systematically and quantitatively compare the response magnitude of each neuron in wakefulness and sleep states across the entire dataset (Figure 5A). PRC neurons showed attenuation with median gain factor of -47% (Confidence Interval CI = [-56%, -34%]) in NREM sleep and -36% (CI= [-43%, -30%]) in REM sleep (Figure 5B). By contrast, AC neurons only exhibited modest changes, with median gain factor of -12% (CI= [-20%, -3%]), and -6% (CI= [-19%, 2%]) in NREM sleep and REM sleep, respectively. The distribution of PRC sleep attenuation gain factors was significantly more negative than that of AC (*p* = 10^−5^ and *p* = 10^−4^ via Mann-Whitney test for NREM sleep, and REM sleep, respectively). Results were largely comparable when gain factors were computed for each stimulus separately (median gain factor in AC: +3% and -11% during NREM and REM sleep respectively; in PRC: -37% and -41% during NREM and REM sleep respectively). Different sound categories (i.e. pure tones, white noise, clicks, vocalizations) were not associated with significantly different attenuation profiles during sleep. Separate analysis of light NREM sleep epochs (Methods) showed a median gain modulation of -26% (CI= [-37%, -17%], not shown).

**Figure 5.**
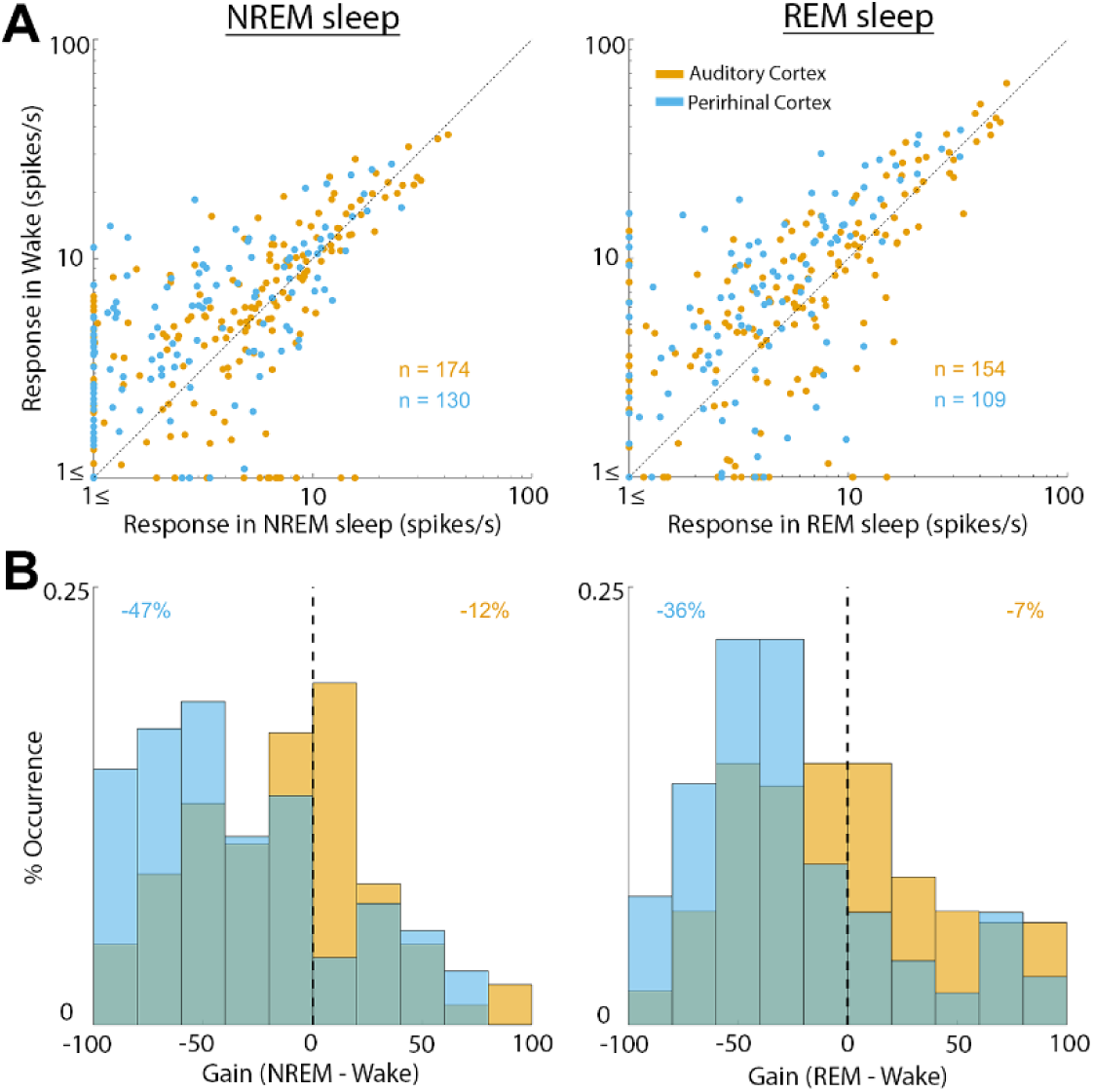
Quantitative comparison of auditory responses in AC and PRC neurons across wakefulness and sleep. (A) Scatter plot of response magnitude in wakefulness (y-axis) vs. NREM sleep (x-axis, left column) or vs. REM sleep (x-axis, right column). Each dot represents one unit and its average response magnitude across all effective stimuli (Methods). Note that AC neurons (orange dots) distribute around the identity line, while PRC neurons (blue dots) are mostly above the diagonal (stronger responses in wakefulness). (B) Distribution of gain factors (Methods) for AC neurons vs. PRC neurons. Colors as above. Vertical dashed line shows zero gain and numbers in upper corners indicate the median gain factor in each neuronal population.

**Figure 6.**
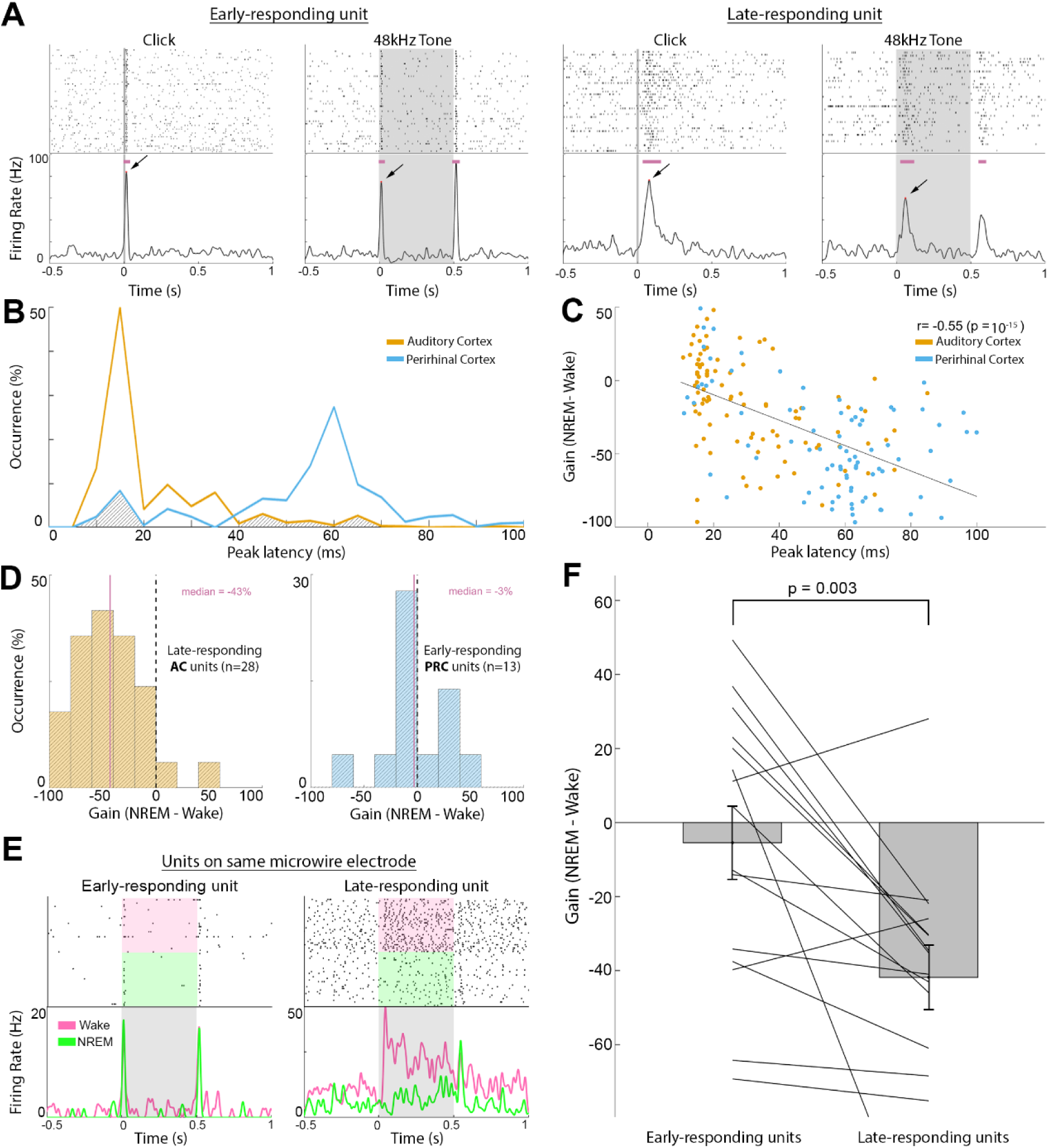
Response latency better predicts attenuation than anatomical region does during NREM sleep. (A) Example responses to click and tone pip stimuli during wakefulness in an early-responding neuron (two left columns) vs. a late-responding neuron (two right columns). Grey shades denote stimulus presentation times, horizontal purple bars indicate automatically-detected response intervals, and arrows mark peak response used for early-vs. late-categorization (latencies from left to right: 16, 14, 80, and 62ms). (B) Distribution of peak response latencies in auditory cortex (orange line, typically <20ms) vs. perirhinal cortex (blue line, typically > 40ms). Gray shading marks clusters with ‘atypical’ latencies in each area (>40ms in AC and <20ms in PRC) that are analyzed separately in panel D. (C) Scatter plot of wake vs. NREM sleep gain factor (y-axis) vs. peak response latency (x-axis) for each neuron, colored accordance to anatomical region. Upper right corner shows Pearson correlation coefficients and corresponding statistical significance, representing a tendency of late-responding neurons to exhibit larger differences in response magnitude between wakefulness and NREM sleep. (D) Examination of neurons with ‘atypical’ latencies for their region (gray shaded areas in panel B). Left, wake/NREM sleep gain factor distribution of AC neurons with long (>40ms) response latency (n=28). Right, wake/NREM sleep gain factor distribution of PRC neurons with short (<20ms) response latency (n=13). Vertical purple lines marks median gain factors (also in purple text on top). Note that latency better predicts sleep attenuation than anatomical region does. (E) Example responses (raster plot & PSTH) of two neuronal clusters identified on the same microelectrode, showing different peak latencies (left, 14ms, right, 45ms) in response to the same sound (3kHz pure tone). Shading indicates stimulus duration. Pink and green colors in raster plot and in PSTH mark trials during wakefulness and NREM sleep, respectively. (F) Sleep attenuation gain factors in 15 pairs of early-responding neurons (left, mean -5.4) and late-responding neurons (right, mean -41) identified on the same microelectrodes. p=0.003 via signed-rank test. Bars and error bars indicate mean and SEM, respectively. Lines connect gain factors in pairs of neurons recorded on the same microelectrodes.

### Response latency of each neuronal cluster is associated with its sleep attenuation

Auditory responsive neuronal clusters fell into two distinct categories based on their peak latency during wakefulness (Figure 6A). Responses of early-responding neurons with peak latencies <20ms were mostly (85%) observed in AC, whereas the majority (89%) of late-responding neurons with peak latencies >40ms were found in PRC (Figure 6B and Methods). Next, we went beyond a binary separation to early- and late-responding neurons, and found a significant correlation (r=-0.55, *p* = 10^−15^) between peak latency and NREM sleep gain attenuation (Figure 6C), and a more modest correlation (r=-0.24, p=0.001) between peak latency and REM sleep gain attenuation (not shown).

Given the tight link between response latency and NREM sleep attenuation, we re-examined how sleep affect the auditory responses of ‘atypical’ neurons (n=41) where latency did not match the typical latency profile of the anatomical region (i.e. late-responding AC neurons or early-responding PRC neurons). We found that late-responding AC neurons show a substantial attenuation profile resembling that observed in PRC (gain factor of -43%, CI= [-60%, -25%]), whereas early-responding PRC neurons only showed modest attenuation in NREM sleep, as was observed in AC (gain factor of -3%, CI= [-16%, 33%], Figure 6D). Finally, in the few cases where two neuronal clusters were identified on the same microwire and showed distinct response latencies (n=15 pairs, Figure 6E), we found a robust (8-fold) difference in gain attenuation during NREM sleep such that early-responding neurons showed a modest average gain (−5.4±9.8%) vs. robust (−41±8.6%) attenuation for late-responding neurons (mean±SEM, p=0.003 via signed-rank test, Figure 6F). In REM sleep, attenuation did not clearly follow response latency as was the case during NREM sleep (not shown), nor did we observe a significant difference between attenuation for neuronal pairs with different latencies recorded on same microelectrodes (p=0.07, +5% in early-responding neurons vs. -20% in late-responding neurons).

## Discussion

The present results reveal how natural sleep differentially affects auditory responses along the cortical sensory hierarchy. We find that PRC exhibit robust attenuation of response magnitude in both NREM sleep and REM sleep, whereas AC neurons are only modestly affected by sleep. Moreover, response attenuation is correlated with response latency in individual neurons. Dividing neurons according to their response latency can explain the NREM sleep attenuation profiles based on cortical region (AC vs. PRC), and can also go beyond anatomical parcellation in showing modest sleep attenuation in early-responding PRC neurons vs. robust attenuation in late-responding AC neurons. Given the preserved auditory responses in previous studies, our results provide the first direct neuronal evidence for attenuation of auditory responses in natural sleep, and pave the road for future studies to dissect the mediating mechanisms.

Downstream to the primary auditory cortex, a previous study in marmosets (Issa and Wang, 2008) showed that secondary auditory cortices also exhibit preserved responses between vigilance states. However, studies of sensory processing beyond early sensory regions during natural sleep are limited. One study (Vinnik et al., 2012) found that a subset of hippocampal neurons in the rat do exhibit robust sensory responses during natural sleep. Another study (Pereira et al., 2007) reported increased hippocampal responses to microsimulation of the rodent somatosensory whisker system during sleep. This discrepancy with the robust attenuation we find here in rat PRC may reflect differences in sensory modalities, or differences between natural stimuli used here vs. ultra-short electrical stimulation.

In AC, comparable early responses in natural sleep observed here are in line with several recent reports (Pena et al., 1999; Edeline et al., 2001; Issa and Wang, 2008; Nir et al., 2013). In addition, a recent study comparing auditory responses at the neuronal level in humans across wakefulness and light anesthesia found a similar distinction between relatively preserved early responses in AC vs. robust attenuation of late responses in association cortex (Krom et al., 2018). What could be the source of apparent discrepancy with earlier studies in other sensory modalities that reported attenuated spiking responses in primary cortices during natural sleep? One possibility is that auditory processing is relatively immune to thalamic gating. Along this line, residual sensory processing during sleep has been demonstrated mainly using acoustic and linguistic stimuli (Bastuji and García-Larrea, 1999; Hennevin et al., 2007). However, a closer examination of the original results (Evarts, 1963) indicates that the response latency of attenuated responses is around 80-100ms, in line with our observation that during NREM sleep, response latency better predicts sleep attenuation than anatomical region (Figure 6). Other classical studies in primary somatosensory and visual cortices (Gücer, 1979; Livingstone and Hubel, 1981) only considered neurons with robust responses during wakefulness and quantified their response profiles in sleep, leaving out potential responses that could be stronger/exclusively present in sleep. Future studies comparing responses between vigilance states should take into consideration the existence of sleep responses that diminish during wakefulness.

We find that early AC responses are comparable across both NREM and REM sleep, whereas PRC responses were attenuated in NREM sleep, and to a lesser extent during REM sleep. We propose that the degree of response attenuation in high-order cortex mirrors the depth of sleep as defined behaviorally. Indeed, in the rat arousal thresholds are higher in NREM sleep (compared with REM sleep) and with increased slow wave activity(Neckelmann and Ursin, 1993; Hayat et al., 2019). Along this line, light NREM sleep is associated with a lower arousal threshold (Emris et al., 2010; Hayat et al., 2019) and we find it exhibits a more modest response attenuation than both NREM sleep and REM sleep.

The robust correlation of NREM sleep attenuation with response latency suggests that NREM sleep may exert a gradual attenuation of cortical responses, rather than imposing a discrete ‘gate’ at a specific brain region. As a first approximation, AC and PRC are associated with different response latency profiles (Lee et al., 2016) and differ also in their degree of response attenuation during sleep. However, a closer examination of neuronal clusters with ‘atypical’ latencies reveals that their attenuation during NREM sleep is better predicted by their classification to early/late-responders rather than their anatomical location (Figure 6). The fact that late responses exhibit stronger attenuation in NREM sleep is in line with the notion of breakdown of cortical effective connectivity during NREM sleep (Massimini et al., 2005) that may lead to greater attenuation with increasing synaptic connections. By contrast, we could not reveal robust relation between response latency and attenuation in REM sleep, in line with the partial recovery of effective connectivity during REM sleep (Massimini et al., 2010). Thus, during REM sleep other mechanisms may be at play. For example, although speculative, it is possible that during REM sleep ongoing mnemonic activity during dreaming interferes with bottom-up sensory processes (Nir and Tononi, 2010; Andrillon et al., 2016).

Of note, we employed a passive auditory stimulation scheme where sounds were not associated with specific behaviors or rewards. This approach allowed us to distill changes related to vigilance state per-se, without confounding processes that boost responses to behaviorally-relevant stimuli during wakefulness. Future studies could compare how sleep may differentially affect responses to such salient stimuli beyond the effects described here.

In conclusion, we find a robust attenuation of late responses in high-order sensory association cortex during natural sleep at the level of individual neuronal spiking. Building on the anatomical and temporal markers of sleep attenuation found here, future studies can elucidate the precise mechanisms underlying the sensory disconnection of sleep.

## Acknowledgements

Supported by the Israel Science Foundation (ISF) grant 1326/15 and 51/11 (I-CORE cognitive sciences) and the Adelis Foundation (YN), by ISF grant 762/16 and the European Society of Anesthesia young investigator start-up grant (AJK), and by an Azrieli Fellowship award from the Azrieli Foundation (YS). We thank Israel Nelken for suggesting to focus on perirhinal cortex as high-order region. Hanna Hayat and Amit Marmelshtein for their input, and all members of Nir lab for discussions.

## References

Allen TA, Furtak SC, Brown TH (2007) Single-unit responses to 22kHz ultrasonic vocalizations in rat perirhinal cortex. Behav Brain Res 182:327–336 Available at: http://linkinghub.elsevier.com/retrieve/pii/S0166432807001477.

Andrillon T, Poulsen AT, Hansen LK, Léger D, Kouider S (2016) Neural Markers of Responsiveness to the Environment in Human Sleep. J Neurosci 36:6583–6596 Available at: http://www.jneurosci.org/lookup/doi/10.1523/JNEUROSCI.0902-16.2016.

Bastuji H, García-Larrea L (1999) Evoked potentials as a tool for the investigation of human sleep. Sleep Med Rev 3:23–45 Available at: http://www.ncbi.nlm.nih.gov/pubmed/15310488.

Benjamini Y, Yekutieli D (2001) The control of the false discovery rate in multiple testing under dependency. Ann Stat 29:1165–1188 Available at: https://projecteuclid.org/euclid.aos/1013699998.

Campbell MJ, Gardner MJ (1988) Calculating confidence intervals for some non-parametric analyses. Available at: https://www.ncbi.nlm.nih.gov/pmc/articles/PMC2545906/pdf/bmj00286-0037.pdf [Accessed March 25, 2019].

Carskadon MA, Dement WC (2011) Normal human sleep: an overview. In: Principles and practice of sleep medicine, 5th ed. (Kryger MH, Roth T, Dement WC, eds), pp 16–26. St. Louis (MO): Saunders/Elsevier.

Cirelli C, Tononi G (2008) Is Sleep Essential? PLoS Biol 6:e216 Available at: http://dx.plos.org/10.1371/journal.pbio.0060216.

Datta S, Hobson J a (2000) The rat as an experimental model for sleep neurophysiology. Behav Neurosci 114:1239–1244 Available at: http://www.ncbi.nlm.nih.gov/pubmed/11142656.

Decurtis M, Pare D (2004) The rhinal cortices: a wall of inhibition between the neocortex and the hippocampus. Prog Neurobiol 74:101–110 Available at: http://linkinghub.elsevier.com/retrieve/pii/S0301008204001455.

Doron NN, Ledoux JE, Semple MN (2002) Redefining the tonotopic core of rat auditory cortex: physiological evidence for a posterior field. J Comp Neurol 453:345–360 Available at: http://www.ncbi.nlm.nih.gov/pubmed/12389207.

Edeline JM, Dutrieux G, Manunta Y, Hennevin E (2001) Diversity of receptive field changes in auditory cortex during natural sleep. Eur J Neurosci 14:1865–1880.

Emris U, Krakow K, Voss U (2010) Arousal thresholds during human tonic and phasic REM sleep. J Sleep Res 19:400–406 Available at: http://doi.wiley.com/10.1111/j.1365-2869.2010.00831.x.

Evarts E V (1963) Photically evoked responses in visual cortex units during sleep and waking. J Neurophysiol Publ 26:229–248.

Furtak SC, Allen TA, Brown TH (2007) Single-Unit Firing in Rat Perirhinal Cortex Caused by Fear Conditioning to Arbitrary and Ecological Stimuli. J Neurosci 27:12277–12291 Available at: http://www.jneurosci.org/cgi/doi/10.1523/JNEUROSCI.1653-07.2007.

Gücer G (1979) The effect of sleep upon the transmission of afferent activity in the somatic afferent system. Exp Brain Res 34:287–298.

Hayat H, Regev N, Matosevich N, Sales A, Paredes-Rodriguez E, Krom AJ, Bergman L, Li Y, Lavigne M, Kremer EJ, Yizhar O, Pickering AE, Nir Y (2019) Locus-coeruleus norepinephrine activity gates sensory-evoked awakenings from sleep. BioArchive.

Hennevin E, Huetz C, Edeline JM (2007) Neural representations during sleep: From sensory processing to memory traces. Neurobiol Learn Mem 87:416–440.

Issa EB, Wang X (2008) Sensory responses during sleep in primate primary and secondary auditory cortex. J Neurosci 28:14467–14480 Available at: http://www.pubmedcentral.nih.gov/articlerender.fcgi?artid=3844765&tool=pmcentrez&rendertype=abstract [Accessed March 24, 2014].

Kakigi R, Naka D, Okusa T, Wang X, Inui K, Qiu Y, Tran TD, Miki K, Tamura Y, Nguyen TB, Watanabe S, Hoshiyama M (2003) Sensory perception during sleep in humans: a magnetoencephalograhic study. Sleep Med 4:493–507 Available at: http://www.ncbi.nlm.nih.gov/pubmed/14607343.

Kealy J, Commins S (2011) The rat perirhinal cortex: A review of anatomy, physiology, plasticity, and function. Prog Neurobiol 93:522–548.

Krom AJ, Marmelshtein A, Gelbard-Sagiv H, Tankus A, Hayat D, Matot I, Strauss I, Fahoum F, Soehle M, Boström J, Mormann F, Fried I, Nir Y (2018) Anesthesia-induced loss of consciousness disrupts auditory responses beyond primary cortex. bioRxiv:502385 Available at: https://www.biorxiv.org/content/10.1101/502385v1 [Accessed March 26, 2019].

Lee CM, Osman AF, Volgushev M, Escabí MA, Read HL (2016) Neural spike-timing patterns vary with sound shape and periodicity in three auditory cortical fields. J Neurophysiol 115:1886–1904 Available at: http://www.physiology.org/doi/10.1152/jn.00784.2015 [Accessed October 11, 2018].

Livingstone MS, Hubel DH (1981) Effects of sleep and arousal on the processing of visual information in the cat. Nature 291:554–561 Available at: http://www.ncbi.nlm.nih.gov/pubmed/6165893.

Makov S, Sharon O, Ding N, Ben-Shachar M, Nir Y, Zion Golumbic E (2017) Sleep Disrupts High-Level Speech Parsing Despite Significant Basic Auditory Processing. J Neurosci 37:7772–7781 Available at: http://www.jneurosci.org/lookup/doi/10.1523/JNEUROSCI.0168-17.2017.

Massimini M, Ferrarelli F, Huber R, Esser SK, Singh H, Tononi G (2005) Breakdown of cortical effective connectivity during sleep. Science 309:2228–2232 Available at: http://www.ncbi.nlm.nih.gov/pubmed/16195466 [Accessed July 10, 2018].

Massimini M, Ferrarelli F, Murphy MJ, Huber R, Riedner BA, Casarotto S, Tononi G (2010) Cortical reactivity and effective connectivity during REM sleep in humans. Cogn Neurosci 1:176–183 Available at: http://www.tandfonline.com/doi/abs/10.1080/17588921003731578.

McCormick DA, Bal T (1994) Sensory gating mechanisms of the thalamus. Curr Opin Neurobiol 4:550–556 Available at: http://linkinghub.elsevier.com/retrieve/pii/0959438894900566 [Accessed February 5, 2015].

Murakami M, Kashiwadani H, Kirino Y, Mori K (2005) State-Dependent Sensory Gating in Olfactory Cortex. Neuron 46:285–296.

Neckelmann D, Ursin R (1993) Sleep stages and EEG power spectrum in relation to acoustical stimulus arousal threshold in the rat. Sleep 16:467–477.

Nir Y, Tononi G (2010) Dreaming and the brain: from phenomenology to neurophysiology. Trends Cogn Sci 14:88–100.

Nir Y, Vyazovskiy V V, Cirelli C, Banks MI, Tononi G (2013) Auditory Responses and Stimulus-Specific Adaptation in Rat Auditory Cortex are Preserved Across NREM and REM Sleep. Cereb Cortex 25:1362–1378 Available at: http://www.ncbi.nlm.nih.gov/pubmed/24323498.

Oudiette D, Paller KA (2013) Upgrading the sleeping brain with targeted memory reactivation. Trends Cogn Sci 17:142–149 Available at: https://linkinghub.elsevier.com/retrieve/pii/S136466131300020X [Accessed March 24, 2019].

Pelletier JG (2004) Low-Probability Transmission of Neocortical and Entorhinal Impulses Through the Perirhinal Cortex. J Neurophysiol 91:2079–2089 Available at: http://jn.physiology.org/cgi/doi/10.1152/jn.01197.2003.

Pena JL, Pérez-Perera L, Bouvier M, Velluti RA (1999) Sleep and wakefulness modulation of the neuronal firing in the auditory cortex of the guinea pig. Brain Res 816:463–470.

Pereira A, Ribeiro S, Wiest M, Moore LC, Pantoja J, Lin S-C, Nicolelis MAL (2007) Processing of tactile information by the hippocampus. Proc Natl Acad Sci 104:18286–18291 Available at: http://www.pnas.org/cgi/doi/10.1073/pnas.0708611104.

Portas CM, Krakow K, Allen P, Josephs O, Armony JL, Frith CD (2000) Auditory Processing across the Sleep-Wake Cycle: Simultaneous EEG and fMRI Monitoring in Humans. Neuron 28:991–999.

Profant O, Burianová J, Syka J (2013) The response properties of neurons in different fields of the auditory cortex in the rat. Hear Res 296:51–59 Available at: http://dx.doi.org/10.1016/j.heares.2012.11.021 [Accessed March 24, 2019].

Quiroga RQ, Nadasdy Z, Ben-Shaul Y (2004) Unsupervised spike detection and sorting with wavelets and superparamagnetic clustering. Neural Comput 16:1661–1687.

Rasch B, Buchel C, Gais S, Born J (2007) Odor Cues During Slow-Wave Sleep Prompt Declarative Memory Consolidation. Science (80-) 315:1426–1429 Available at: http://www.ncbi.nlm.nih.gov/pubmed/17347444 [Accessed March 24, 2019].

Rothschild G, Eban E, Frank LM (2017) A cortical-hippocampal-cortical loop of information processing during memory consolidation. Nat Neurosci 20:251–259 Available at: http://www.nature.com/articles/nn.4457 [Accessed March 24, 2019].

Sela Y, Vyazovskiy V V, Cirelli C, Tononi G, Nir Y (2016) Responses in Rat Core Auditory Cortex are Preserved during Sleep Spindle Oscillations. Sleep Available at: http://www.ncbi.nlm.nih.gov/pubmed/26856904.

Steriade M (2003) Neuronal substrate of sleep and epilepsy. Cambridge (England): Cambridge University Press.

Steriade M, McCarley RW (1990) Brainstem Control of Wakefulness and Sleep. Boston, MA: Springer US. Available at: http://link.springer.com/10.1007/978-1-4757-4669-3 [Accessed August 26, 2015].

Strauss M, Sitt JD, King J-R, Elbaz M, Azizi L, Buiatti M, Naccache L, van Wassenhove V, Dehaene S (2015) Disruption of hierarchical predictive coding during sleep. Proc Natl Acad Sci 112:E1353–E1362 Available at: http://www.pnas.org/cgi/doi/10.1073/pnas.1501026112 [Accessed February 10, 2019].

Vinnik E, Antopolskiy S, Itskov PM, Diamond ME (2012) Auditory stimuli elicit hippocampal neuronal responses during sleep. Front Syst Neurosci 6:49 Available at: http://journal.frontiersin.org/article/10.3389/fnsys.2012.00049/abstract.

